# Sorting of molecular shuttles by designing electrical and mechanical properties of microtubules

**DOI:** 10.1101/107458

**Authors:** Naoto Isozaki, Hirofumi Shintaku, Hidetoshi Kotera, Taviare L. Hawkins, Jennifer L. Ross, Ryuji Yokokawa

**Affiliations:** Department of Micro Engineering, Kyoto University, Kyoto 615-8540, Japan; Department of Physics, University of Wisconsin-La Crosse, La Crosse, Wisconsin 54601, USA; Department of Physics, University of Massachusetts–Amherst, Amherst, Massachusetts 01003, USA

**Author notes:** To whom correspondence should be addressed: Department of Micro Engineering, Kyoto University, Kyoto Daigaku-Katsura, Nishikyo-ku, Kyoto 615-8540, Japan.

## Abstract

Kinesin-driven microtubules have been a focus to serve as molecular shuttles to replace multiple on-chip functions in micro total analysis systems μTAS). Although transport, concentration, and detection of target molecules have been demonstrated, controllability of transport directions is still a major challenge. To define multiple moving directions for selective molecular transport, we integrated the bottom-up molecular design of microtubules and the top-down design of a microfluidic device. The surface charge density and stiffness of microtubules were controlled, allowing us to create three different types of microtubules with different gliding directions corresponding to their electrical and mechanical properties. The measured curvature of gliding microtubules enabled us to optimize the size and design of the device for molecular sorting in a top-down approach. The integrated bottom-up and top-down design achieved separation of stiff microtubules from negatively-charged soft microtubules with approximately 80% efficiency under an electric field. Our method is the first to sort multiple microtubules by integrating molecular control and microfluidic device design, and is applicable to multiplexed molecular sorters.

---

Kinesin motor proteins and microtubule (MT) cytoskeletal filaments show promise as in vitro nano-scale actuator platform for nanobiotechnology applications. MTs are rigid, polar, dynamic cytoskeletal filaments that serve as mechanically supportive cellular structures. MTs also serve as the highway system for intracellular transport by kinesin and dynein motor proteins. Such motor-driven active transport uses the hydrolysis of adenosine triphosphate (ATP) and is indispensable for maintaining cellular functions.^1,2^ In the cell, kinesin motor proteins walk along MT s to deliver vesicular and small molecule cargos to destinations in an in vivo viscous environment. The MT-kinesin transport system can be reconstituted in vitro for cargo transport or inverted to have molecular motors transport MTs.

The MT-kinesin system can be combined with microfluidics, to enable simultaneous control of aqueous, bulk chemical composition with direct transport of molecular-scale cargos. These systems promise unprecedented control over nanoscale delivery—not just bulk flow and chemical reactions, and has matured into a micro total analysis systems μTAS) that could replace on-chip functions in μTAS.^3–5^ These μTAS systems enable a wide range of applications using motor-driven MTs as molecular shuttles such as molecular transporters, sorters, concentrators, and detectors. MT-kinesin μTAS load cargos onto gliding MTs via avidin-biotin binding,^5,6^ antigen-antibody interaction,^7,8^ or DNA hybridization^9,10^ to be sorted, transported, and detected on the chip. Although individual functions have been demonstrated, a long-lasting challenge that hampers practical usage of MT-kinesin is the directional control of gliding MT shuttles, because they glide in random directions due to Brownian motion acting on the free leading tip (minus ends). MT trajectories can be controlled by photoresist tracks,^5,11,12^ fluid shear flow,^13,14^ magnetic fields,^15^ and electric fields.^16–18^ However, these methods limit the control of MT transport to only one destination; without an active control, all MT shuttles behave in the same manner and glide toward the same destination.^16^ Here, we tackled this challenge of multiple-cargo sorting by integrating a bottom-up molecular design of MTs and top-down design of the microfluidic device. Using these methods, we demonstrate, for the first time, MT gliding to two destinations as a highly-efficient MT sorter under a single external field.

For the bottom-up manipulation of MTs, we focused on two MT properties to be designed: electrical and mechanical properties. Our group has already reported that the surface charge density of MTs is negatively correlated with the radius of the trajectory curvature.^19^ Design of the MT surface charge density enabled us to guide MTs toward multiple directions under a given electric field. Here, we additionally manipulate the mechanical stiffness of the MTs measured as flexural rigidity (*κ*) or persistence length (*L*_p_). In vivo, MTs control their stiffness depending on the intracellular roles of the filament, i.e. stiffer MTs are needed in the axon to support its long structure, whereas more flexible MTs are needed in a proliferating cell to enable rapid redistribution.^20^ Many factors for altering MT stiffness have been reported including: MT stabilizing agents, nucleotides, MT-associated proteins (MAPs), and growth rates. However, there are still controversies on stiffening and softening factors, and measured stiffness has never before been used to manipulate MT gliding directions.^17, 20–38^ The cantilever-beam model suggests that MT stiffness is proportional to the persistence of the gliding trajectory,^19^ so we proposed designing both electrical and mechanical properties simultaneously to improve controllability of the gliding directions. We predicted that MTs with low surface charge density and high stiffness will have larger radii of curvature trajectories (longer persistence) than those with high surface charge density and low stiffness (Fig. 1).

**Figure 1.**
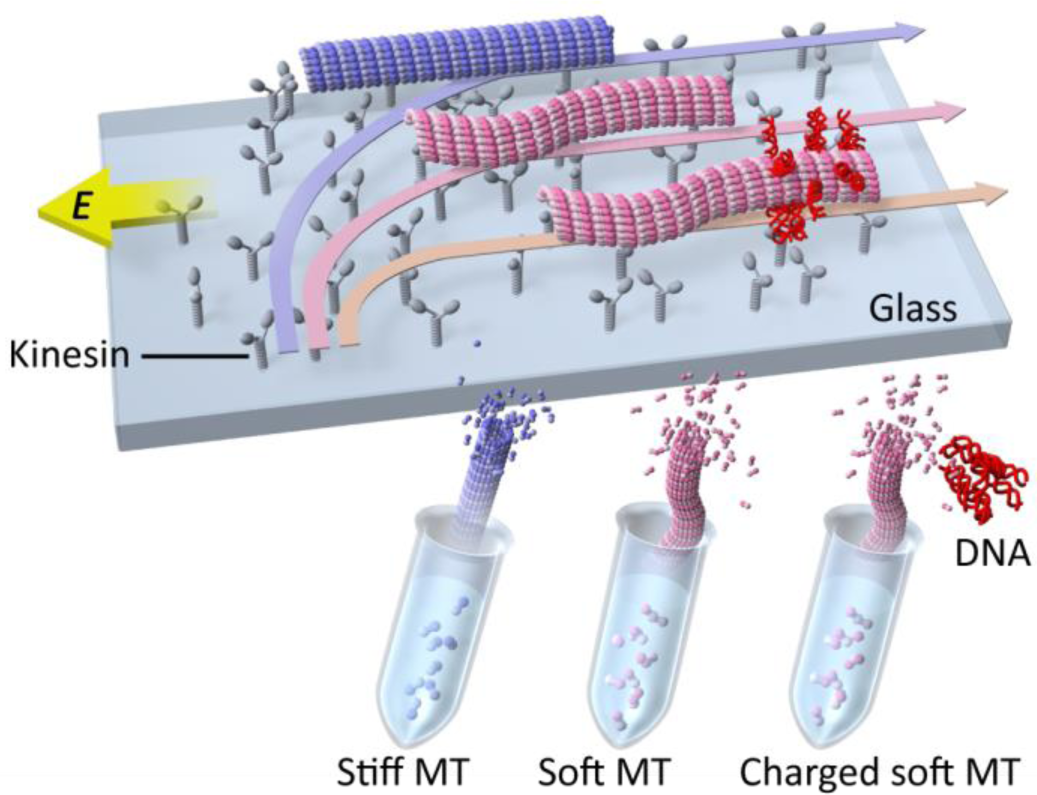
Schematic representation of MT sorting under a given electric field. Designing MT properties controls the trajectories of kinesin-propelled MTs. Various types of MTs were polymerized under different conditions in microtubes. When an electric field, *E*, is applied perpendicular to gliding MTs, their gliding directions are gradually oriented toward the anode. MTs are transported toward different destinations corresponding to their stiffness and surface charge density.

In this study, we first investigated the stiffness of MTs to find the highest and lowest rigidity under various polymerizing conditions—using nucleotides, growth rate, and the neuronal MAP, tau^25^ The *L*_p_ of the designed MTs was measured using thermal fluctuations, and the relationship of *L*_p_ to the gliding trajectories under an electric field was obtained. We further modified the electrical properties of MTs by DNA labelling filaments. We have shown that adding DNA can alter the radius of curvature by 2.3- fold.^19^ In addition to the bottom-up designs of MT electrical and mechanical properties, we took a top-down approach to design a microfluidic device to separate MTs according to measured radii of trajectory curvatures (Fig. 1). This integrated approach enabled us to guide two types of MTs toward two different destinations and separate them in the device corresponding to their properties, resulting in MT sorting at approximately 80% efficiency. Our work is the first to actualize differential sorting of molecular shuttles to multiple destinations. This work is useful for a multiplex molecular sorter in a microfluidic device.

## Results

### Variable persistence length under different polymerizing conditions

We investigated five types of MTs listed in Table 1: Tau-bound slowly-polymerized guanylyl (α,β)methylenediphosphonate GMPCPP-MT (MT-1), tau-free slowly-polymerized GMPCPP-MT (MT-2), tau-bound slowly-polymerized GTP-MT (MT-3), tau-free slowly-polymerized GTP-MT (MT-4), and tau-free quickly-polymerized GTP-MT (MT-5). Each MT type was elongated as segments from short MT seeds made in the presence of GMPCPP with tubulin dimers that were biotinylated to immobilize them onto a glass substrate to measure their *L*_p_. The growth rate in the elongation process was controlled by using either 10 μM or 30 μM free tubulin in the presence of GTP during polymerization (Fig. 2a). To measure the length of the elongated segments, MTs were sampled 0.5-60 min after elongation started, and all MTs were stabilized by paclitaxel just after sampling.

**Table 1.** The five MTs investigated.

|  |  |  |  |  |  |
| --- | --- | --- | --- | --- | --- |
| Name | MT-1 | MT-2 | MT-3 | MT-4 | MT-5 |
| Growth rate | Slow |  |  |  | Fast |
| Nucleotide | GMPCPP |  | GTP |  |  |
| Tau binding | Yes | No | Yes | No |  |

**Figure 2.**
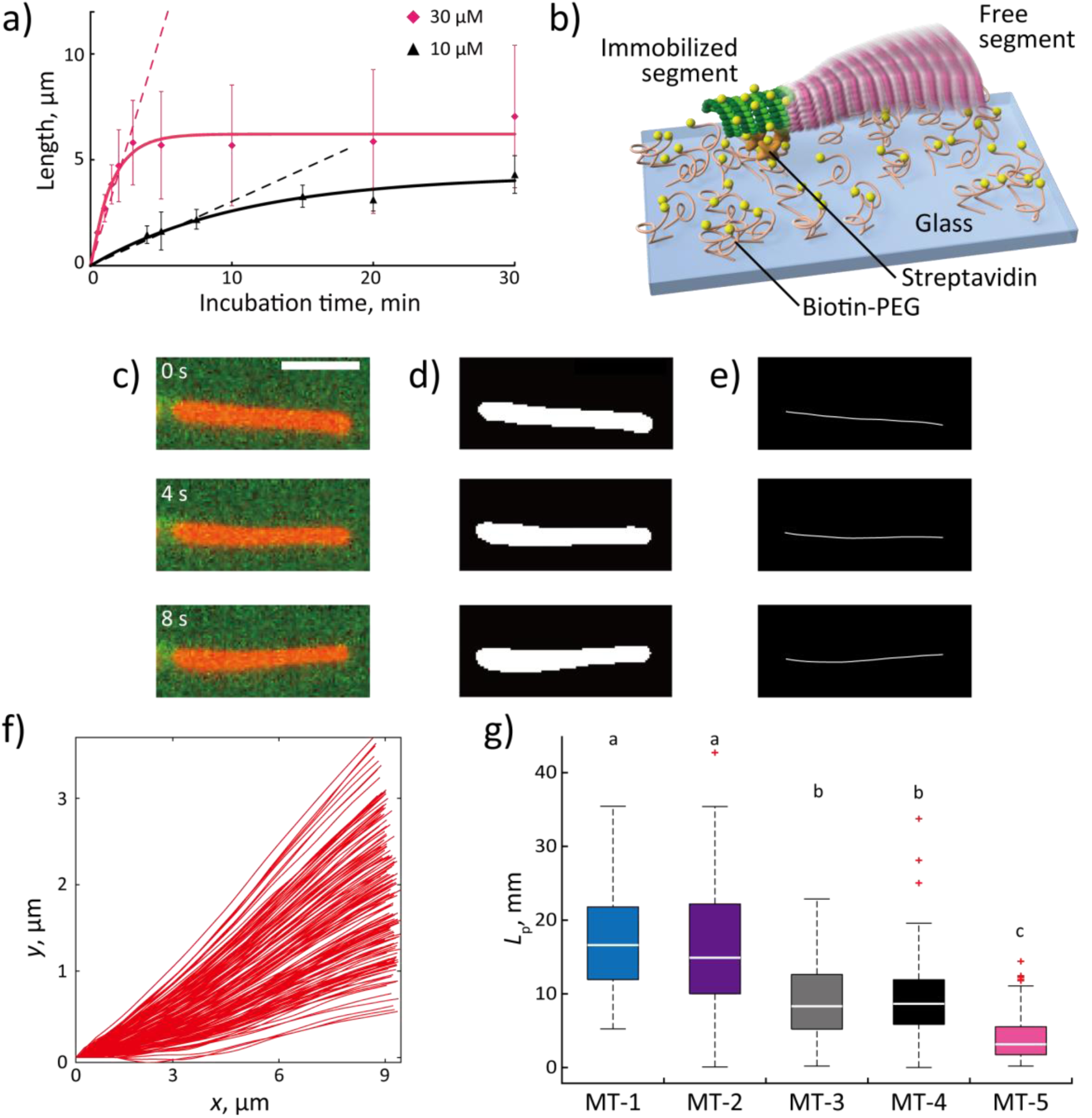
Control of MT persistence length by polymerizing conditions. (**a**) Time course of MT length in the presence of 30 μM (red diamonds) and 10 μM tubulin concentrations (black triangles). Mean ± SD are shown and least-squares fitting to an exponential curve (solid lines). Dashed lines show least-squares fitting of lines for the first several minutes (*R*^2^ > 0.92). (**b**) The schematic representation of the measurement system. MTs were partially immobilized onto a biotin-coated substrate by streptavidin and biotinylated MT seeds. (**c**) Sequential images of a fluctuating MT (red). The left segment (light green) was immobilized. Scale bar = 5 μm. Images were (**d**) binarized and (**e**) skeletonized with the FIESTA software. (**f**) Superposition of the whole shape of a fluctuating MT for each frame. (**g**) Box plots of *L*_p_ for MT-1 (*n* = 114 measurements), MT-2 (*n* = 128), MT-3 (*n* = 100), MT-4 (*n* = 86), and MT-5 (*n* = 126). Red plus signs mean outliers defined by 1.5 interquartile ranges. There are no significant differences between identical lowercase letters by the Steel-Dwass test at a critical value of *p* < 0.01.

To determine the time course against the lengths of the elongated segments, the distribution of MT length was plotted and fit to Gaussian functions to find the mean and standard deviation (Supplementary Fig. 1). Only normally distributed data were used and plotted (Fig. 2a). MTs elongated linearly in the first several minutes and then reached plateaus, as expected. Data were fitted as solid lines by an exponential decay, *L*(*t*) = *L*_max_(1-exp(-*t*/*τ*)). Here, *L* is the MT length over time, *t*, *L*_max_ is the upper limit of *L*, and *τ* is the characteristic time constant. *τ* and *L*_max_ were 1.53 min and 6.22 μm for 30 μM tubulin (*R*^2^ = 0.94), and 10.8 min and 4.28 μm for 10 μM tubulin (*R*^2^ = 0.94), respectively. Growth rates defined by the slopes at early times (dashed lines) were calculated as 2.19 ± 0.15 μm min^−1^ for 30 μM tubulin (*R*^2^ = 0.92) and 0.303 ± 0.017 μm min^-1^ for 10 μM tubulin (*R*^2^ = 0.93). These two concentrations successfully controlled the growth rate, and MTs elongated faster at higher tubulin concentrations, as expected. The measured growth rates are within the range of previous in vitro experiments (0.135-2.56 μm min^-1^).^39–45^

The *L*_p_ was measured for MT-1 to MT-5 (Figs. 2b–g). The biotinylated seed was immobilized onto a glass substrate via biotin-streptavidin and the elongated segment was free to fluctuate under thermal driving (Figs. 2b,c and Supplementary Movie 1). Fluorescent images of MTs were converted to binary images and skeletonized via Gaussian fitting using MT tracking software, FIESTA (Figs. 2c–f)^46^ *L*_p_ was derived by equating thermal energy with bending energy of the MTs, on the assumption that a freely fluctuating segment behaved as a cantilever beam clamped at the immobilized biotinylated segment (see Supporting Information and Supplementary Fig. 2 for details). For each type of MT the distribution of the *L*_p_ was measured and shown to have a normal distribution (Supplementary Fig. 3). MTs were categorised into three groups according to *L*_p_ (Fig. 2g): stiff MTs (MT-1 and MT-2), soft MT (MT-5), and intermediate (MT-3 and MT-4). From our data, we see no significant differences between MT-1 and MT-2, or MT-3 and MT-4 implying that tau binding did not affect *L*_p_ of either GTP-or GWPCPP-polymerized MTs when MTs were slowly polymerized. The nucleotide had a substantial effect on the *L*_p_, since significant differences were found between MT-1 and MT-3, or MT-2 and MT-4. From our work, we observe that *L*_p_ increased approximately two-fold when GTP was replaced by GWPCPP at 10 μM tubulin. Finally, the dependence of *L*_p_ on the growth rate was significant, as demonstrated by the difference between MT-4 and MT-5: *L*_p_ decreased approximately 2.5-fold with the increase of growth rate from 0.303 μm min^−1^ to 2.19 μm min^−1^. Therefore, we obtained three MT groups with different *L*_p_ values by changing the nucleotide and growth rate during polymerization. In addition, the length of the fluctuating segment ranged from 2 μm to 17 μm, and we found no correlation between the filament contour length and *L*_p_ (Supplementary Fig. 4).

### Dependence of radius of trajectory curvature on MT properties

When an electric field is applied in the negative *x*-direction of the *xy*-plane and the MTs enter the field at the origin in the positive *y*-direction, the MT trajectory is defined as follows:

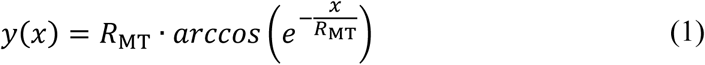

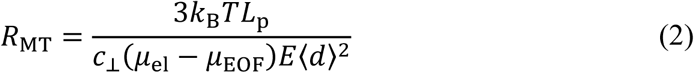

where *k*_B_ is the Boltzmann constant, *T* is the temperature, c_⊥_ is the perpendicular Stokes drag coefficient per unit length of an MT tethered to the surface via kinesin, *μ*_el_ is electrophoretic mobility, *μ*_EOF_ is electroosmotic mobility, *E* is the field intensity, and *<d>* is the average distance between kinesin molecules.^16,17^ The electric field biases MT gliding directions toward the anode with the curvature parameter, *R*_MT_, corresponding to the radius of the curvature. MT-1 and MT-2 had the largest *L*_p_, and MT-5 had the smallest as shown in Fig. 2g. This allowed us to select MT-1 and MT-2 for large *R*_MT_, and MT-5 for small *R*_MT_. To further decrease *R*_MT_ by increasing the *μ*_el_ of MTs, MT-5 was selected for labelling with 50-bp DNA.^19^ Thus, three MT groups with different *R*_MT_ were prepared; MT-1 or MT-2 (stiff MT, large *R*_MT_), MT-5 (soft MT, middle *R*_MT_), and DNA-labelled MT-5 (negatively-charged soft MT, small *R*_MT_). Types MT-1, MT-2 and MT-5 were polymerized under the designated conditions as uniform segments without seed MTs. DNA-labelled MT-5 consisted of a short biotinylated minus-end and a non-labelled plus-end segment. This design was employed to prevent the steric hindrance that disturbs MT gliding if the entire MT surface is coated with streptavidin and DNA molecules.^47^ Moreover, the short minus-end segment was enough to be labelled with DNA, because the gliding direction depends on the properties of the MT leading tips. Hereafter, the partially DNA-labelled MT is referred to as MT-5’.

These four MT types were assayed in a microfluidic channel to directly measure the **A**MT under an electric field (Fig. 3a). The channel was fabricated with polydimethylsiloxane (PDMS) to have a 10 *μ*m height. After nonspecific adsorption of α-casein and kinesin to the PDMS channel, MT gliding was initiated by 0.5 mM ATP. An applied electric field of 5.3 kV m^−1^ guided the MTs toward the anode and *R*_MT_, as defined by equation (1), was measured (Fig. 3b).

**Figure 3.**
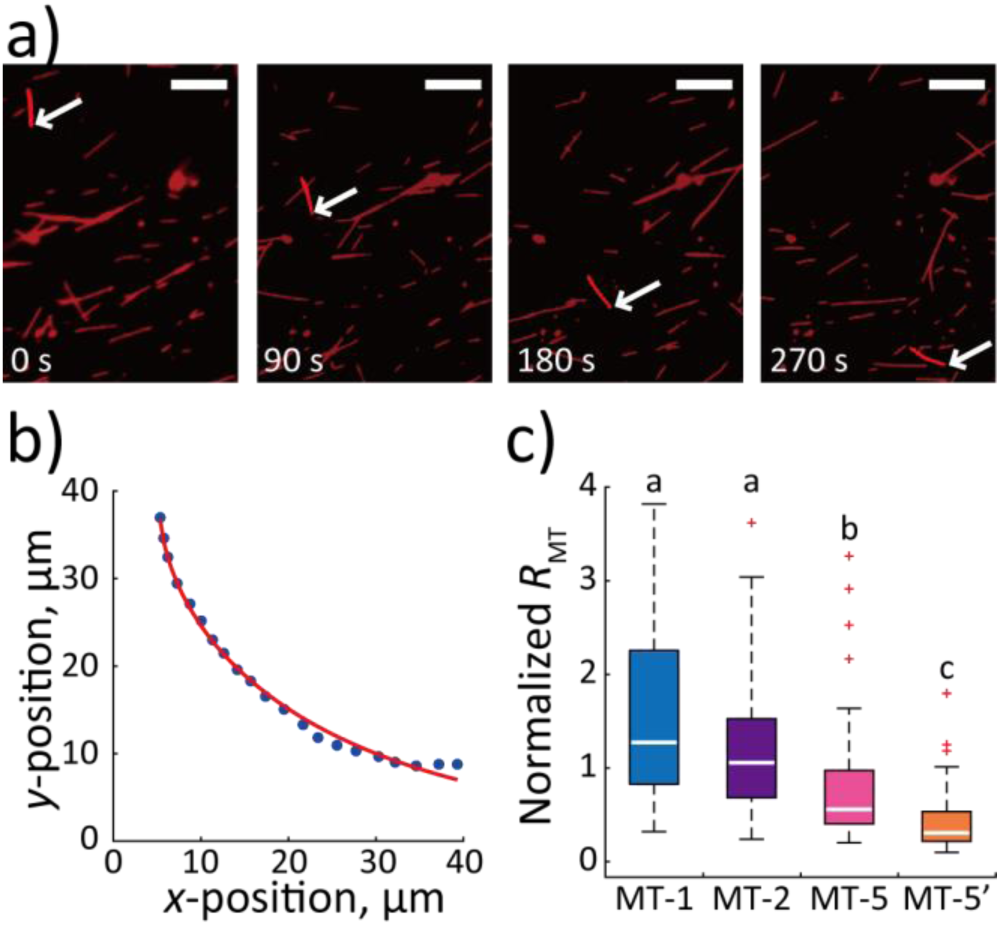
Control of MT gliding directions via the bottom-up design of MT electrical and mechanical properties. (**a**) Sequential images of a gliding MT-2 (highlighted). An electric field of 5.3 kV m^-1^ was applied from the right-to left-hand side of images. White arrows indicate the leading tip of the MT. Scale bar = 10 μm. (**b**) Trajectory and fitted result for the MT shown in (**a**). The leading tip was tracked (blue dots) and fitted with equation (1) as a red line (*R*^2^ = 0.99) to obtain *R*_MT_. (**c**) Box plots of normalized *R*_MT_ for MT-1 (*n* = 32), MT-2 (*n* = 141), MT-5 (*n* = 62), and MT-5’ (n = 62). **A**MT were normalized to mean *R*_MT_ for MT-2. Red plus signs mean outliers defined by 1.5 interquartile ranges. No significant differences were observed between MT-1 and MT-2 (a). MT-5 (b) and MT-5’ (c) showed significant differences with (a) and with each other as determined by the Steel-Dwass test at a critical value of *p* < 0.01. MT trajectories followed equation (1) with *R*^2^ > 0.98.

We measured the normalized *R*_MT_ values for the four MT types (Fig. 3c), which could be categorised into three groups; MTs with large (a: MT-1 and MT-2), middle (b: MT-5), and small (c: MT-5’) *R*_MT_ (Supplementary Fig. 5 shows the raw data). The insignificant difference between MT-1 and MT-2, and the significant difference between MT-1 and MT-2, and MT-5 reflect the differences in *L*_p_ values as shown in Fig. 2g. This positive correlation between *R*_MT_ and *L*_p_ follows equation (2). In addition, MT-5’ showed significantly smaller *R*_MT_ than MT-5. The increase of *μ*_el_ of the leading tips via DNA labelling decreased the *R*_MT_, which also followed equation (2). Therefore, *R*_MT_ was controlled by designing *L*_p_ and *μ*_el_; larger *Lp* and smaller *μ*_el_ produced large *R*_MT_. In order to test sorting, we used either MT-2 and MT-5 or MT-2 and MT-5’.

### Device design and MT sorting

The microfluidic device was designed based on the measured *R*_MT_ values: 40.3 μm (*n* = 90) for MT-2, 21.0 μm (*n* = 57) for MT-5, and 14.5 μm (*n* = 39) for MT-5’ under 5.3 kV m^−1^. Here, three points were considered to define the dimensions of the device: (1) the separation channel needs to be <1 mm width to obtain a uniform electric field;^48^ (2) the cross-contamination should be decreased by increasing the difference of *R*_MT_ to be used as a molecular sorter; (3) to observe MT separation in a field of view, the travel distance in the *y*-direction of either MT group should be less than 80 μm. As equation (2) suggests the difference in *R*_MT_ is inversely proportional to the field intensity, we recalculated *R*_MT_ with a field intensity of 3 kV m^-1^, and the MT trajectories were predicted with equation (1); travelling distances in the *y-* direction were 84.6 μm for MT-2, 58.8 μm for MT-5, and 35.2 μm for MT-5’ at *x* = 70 μm (Supplementary Fig. 6). Therefore, we set the separation wall at (*x*, *y*) = (≥70 μm, 70 μm) to demonstrate the separation of MT-2 from MT-5, and MT-2 from MT-5’ using the same device.

The device can be divided into three areas: MT landing area, MT alignment area, and MT sorting area with the MT separation wall (Figs. 4a-c). MT s landed only on the landing area surface due to the flow generated by pressure differences between reservoirs C and D. Filament gliding directions were aligned to the *y*-axis in the alignment area after ATP was introduced. An electric field subsequently rectified the direction of the MTs toward the anode with a different *R*_MT_ reflecting their Lp and *μ*_el_ in the sorting area.

**Figure 4.**
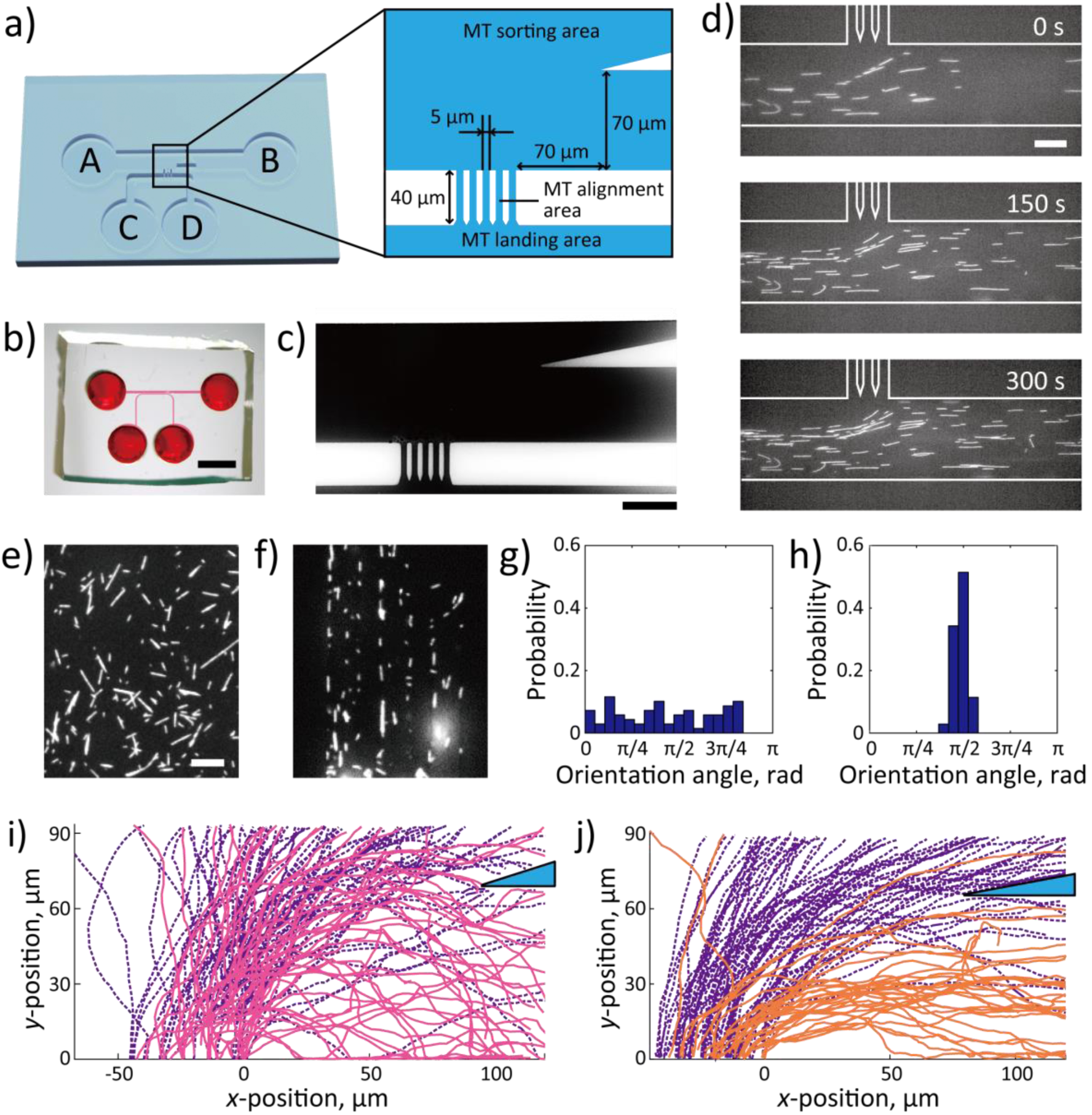
Demonstration of MT sorting by integrating the top-down design of PDMS device with the bottom-up design of MT properties. (**a**) Schematic representation of the PDMS device. The device consisted of three areas: MT landing area (between reservoirs C and D), MT sorting area (between reservoirs A and B), and MT alignment area (between MT landing and sorting areas). Channels in the alignment area were 40 μm in length and 5 μm in width. The channel height was 10 μm. (**b**) Image of the fabricated PDMS device. Red ink indicates the channels. Scale bar = 3 mm. (**c**) Bright field image of the cross section of MT landing, MT alignment, and MT sorting areas. Channels are filled with ink without leakage. Scale bar = 50 μm. (**d**) Sequential images of MT immobilization in the MT landing area. Flow generated by pressure differences among reservoirs prevented MTs from escaping to MT alignment area. White solid lines represent the wall. Scale bar = 20 μm. Fluorescent images of MTs in (**e**) MT landing area and (**f**) MT alignment area. Scale bar = 20 μm. Histograms of MT orientation angles in (**g**) MT landing (*n* = 70) and (**h**) MT alignment areas (*n* = 35). Probabilities are represented by blue bars. MT gliding directions were aligned in the MT alignment area with significant decreases in the standard deviations of the orientation angles. Bin width = π/19 rad. (**i**) MT-2 (purple dashed lines) and MT-5 (red solid lines) were sorted with 57.7% efficiency (*n* = 129). (**j**) MT-2 (purple dashed lines) and MT-5’ (orange solid lines) were sorted with 80.4% efficiency (*n* = 159). Blue triangles represent the PDMS separation wall. An electric field of 3 kV m^-1^ was applied in the negative x-direction. The upper right corners of the MT alignment area are set to origin.

Figure 4d shows the sequential images for MT immobilization in the landing area. The number of landing MTs increased with time and no MT was observed in any other areas. MT alignment was evaluated by comparing the standard deviation of the MT orientation angles in the alignment area (Figs. 4e,g) and in the landing area (Figs. 4f,h). The orientation angle was defined as the angle between the *x*-axis and the measured MT gliding direction. Orientation angles in the alignment area (0.49 ± 0.04 rad, *n* = 35, mean ± SD) showed significantly smaller standard deviations than those in the landing area (0.44 ± 0.27 rad, *n* = 70). Therefore, the MT gliding directions were oriented to the *y*-axis positive direction in the alignment area.

We tracked trajectories in the sorting area for MT-2 and MT-5 (Fig. 4i and Supplementary Movie 2), or MT-2 and MT-5’ (Fig. 4j and Supplementary Movie 3). The sorting efficiency was evaluated by counting the number of MTs separated by the wall. When MT-2 and MT-5 were tested together (*n* = 129), 84.4% of MT-2 glided above and 57.7% of MT-5 glided below the wall. When MT-2 and MT-5’ were tested together (n = 159 in four experiments), 80.4% of MT-2 and 90.4% of MT-5’ were guided above and below the wall. By comparing these results, we demonstrated two significant advances in MT sorting: The sorting efficiency reached about 60% by designing *L*_p_ of MT under different polymerizing conditions, and the efficiency was significantly improved to about 80% by the additive effects of *μ*_el_ to *L*_p_.

## Discussion

In order to create a device that sorts MT shuttles, we modified the MTs themselves. One parameter was to specify the *L*_p_, which we controlled through growth rate, nucleotide content, and the presence of tau protein.

First, we tested the growth rate using two different tubulin concentrations (10 and 30 μM). The lengths of the MTs reached plateaus due to the non-covalent polymers being in dynamic equilibrium with the background concentration of free tubulin (Fig. 2a). As MTs polymerize, the number of free tubulin molecules decreases and some MTs transition from polymerization to catastrophe, which releases tubulin from the filament. Thereafter, tubulin molecules are supplied by catastrophes and individual MTs can grow and shrink. Although the periodic transition between catastrophes and rescues, termed dynamic instability, increased the variation of MT length, the mean MT length becomes constant.^39^ The large standard deviation of the MT length measurements (Fig. 2a) is due to the broad distribution of MT length in Supplementary Fig. 1 in the steady-state phase due to MT dynamic instability.

We found a significant difference in *L*_p_ resulting from different growth rates between MT-4 (slow) and MT-5 (fast) that agreed with previous reports that higher growth rates produced softer MTs.^27,49^ Lp was measured as 3.2 mm for quickly-polymerized MT-5 with a growth rate of 2.19 μm min^−1^ at a tubulin concentration of 30 μM. This result is surprisingly consistent with the *L*_p_ ∼ 3.4 mm (within uncertainty) reported previously for MTs polymerized at 2.40 μm min^−1^ under the tubulin concentration of 28 μM.^49^ Lp was 8.7 mm for the slowly-polymerized MT-4 (growth rate of 0.303 μm min^−1^). It is not surprising that such a low growth rate produced higher *L*_p_ than 6.6 mm previously measured for the lowest reported growth rate (1.50 μm min^−1^).^27^ Recently, higher tubulin concentrations (i.e., higher growth rates) were shown to induce more defects in the MT lattice structure, resulting in lower *L*_p_.^50^ Our work verifies that the control of *L*_p_ by growth rate can lead to the formation of defects.

The difference in nucleotides between MT-2 (GMPCPP) and MT-4 (GTP) agrees with a common understanding: GMPCPP produces stiffer MTs than GTP due to the conformational change in the tubulin dimers, which is a fact that is supported by all prior experiments ^24,30,31,51–56^ However, we found that tau proteins did not show any effect on *L*_p_ in our study, different from previous studies. Hawkins *et al.* reported that copolymerizing tau at 1:100 (tau:tubulin) increased *L*_p_ (0.6 mm to 4 mm), and Felgner *et al.* reported that adding tau to polymerized MTs increased *L*_p_ with the increase of tau (0.92 mm to 2.5 mm).^25,31^ This inconsistency is likely due to the growth rate differences between our work and prior work. All our data with tau present was polymerized very slowly and each type (MT-1,2,3,4) showed higher *L*_p_ (8.3–17 mm) likely due to this very low growth rate. Thus, these MTs may be insensitive to tau proteins due to their lack of defects. Previously, Hawkins et al. tested the effects of paclitaxel, tau, and nucleotide on the stiffness of MTs that were polymerized quickly and found that the order of the addition of various stabilizers affected the outcome.^25^ In the current case, we polymerized with tau, but very slowly. Thus, we conclude that the rate of polymerization, and likely the lack of filament defects, is a more effective regulator of MT stiffness.^23,57^

The dependency of *L*_p_ on MT contour length is more controversial. Our results showed that *L*_p_ is independent of the contour length (Supplementary Fig. 4). Prior groups have measured *L*_p_ for MTs affixed at one end and cantilevered, as we measured, and reported an increase of *L*_p_ with the increase of contour length ^26,30,36,58^ However, others have shown scattered values with a measured contour length for both affixed MTs^25,27,59^ and freely fluctuating MTs.^31,34,35,53,60,61^ More recently, Zhang et *al.* argued that the dependency on length for affixed MTs is negligible for contour lengths >2 μm by numerical simulation.^62^ This supports our results because we only used MTs longer than 2 μm for our *L*_p_ measurements. Our results contribute new understanding of the effects of growth rate on *L*_p_. More importantly, we could consistently measure and modulate *L*_p_ by the growth rate and nucleotide state to alter *R*_MT_ and enable MT sorting.

Our experimental setup for the measurement of *L*_p_ has two advantages from designing two segments. The biotinylated segment was affixed to the glass substrate while the other non-labelled segment fluctuated in a solution. Affixation prevents MT rotation around the longitudinal axis, which could conflate thermally-activated bending with the rotation of a bent filament.^27,34^ Further, by labelling different fluorophores in our segments, we defined the affixed segment in the image analysis which enabled us to precisely find the clamped end of a fluctuating MT, as proposed in a previous study.^27^

Assuming the error introduced from locating the clamped end during the image analyse was minimised, larger errors stem from the MT fluctuating perpendicular (z-direction) to the surface (*xy*-plane) due to the height of the flow cell (10 μm). We observed the projection of MTs onto the *xy*-plane and might have underestimated MT deflection, leading to an overestimation of *L*_p_. Fluctuations in the *z*-direction add an uncertainty to *L*_p_ of up to 8% for the following reasons. The depth of field was 400 nm in our observation system, which means the largest deflection detectable in the z-direction was ∼400 nm. Since MT deflection in the *xy*-plane was 1−3 μm, when the MT tip was elevated 400 nm from the *xy*-plane, the actual deflection is from 1.08 μm (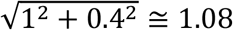) to 3.02 μm (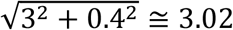). Therefore, the maximum uncertainty of deflection is 8% and the overestimation of *L*_p_ is less than 8%.

We have previously measured the characteristic parameters of MTs as follows: *c*_⊥_ = 1.39 ⊥ 10^−2^ kg m^−1^ s^−1^, *μ*_EOF_ = 1.33 ⊥ 10^−8^ m^2^ V^−1^ s^−1^, *μ*_el_ = 2.03 ⊥ 10^−8^ m^2^ V^−1^ s^−1^ for Alexa Fluor 488-labelled MTs, 2.09 ⊥ 10^−8^ m^2^ V^−1^ s^−1^ for rhodamine-labelled MTs, and 3.02 ⊥ 10^−8^ m^2^ V^−1^ s^−1^ for 50-bp DNA-and rhodamine-labelled MTs.^19^ By substituting these values and the measured RMT into equation 2, we calculated the spacing between kinesin motors <*d*> as 3.0 μm for MT-2, 1.8 μm for MT-5, and 1.5 μm for MT-5’. Our *(d)* are much larger than the distances between kinesin molecules measured in most previous reports: 0.16 μm measured for 0.2 mg ml^−1^ kinesin concentration,^63^ 0.10 μm for 0.07 mg ml^−1,17^ and 0.4 μm for 1.0 μg ml^−1^.^14^ Only one group reported a larger spacing of about 4 μm using a MT-bound microbead manipulated by optical tweezers.^64,65^ Further, why would the distance between kinesins depend on the MT *L*_p_?

There are three possible explanations why <*d*> calculated from the measured *L*_p_ and *R*_MT_ was larger than the distance between kinesins. The first reason could be due to the flexibility of the stalk region of kinesin. Kinesins bound to the MT tips could be extended until detachment, when the electric field was applied to bend the MT. In such a situation, the foremost kinesin is not at the clamped end and the free tip length, deformed by an electric field, becomes larger than the actual distance between kinesins.^14^ This has also been confirmed by a numerical simulation using the same cantilever model by Nitta *et al.* (personal communication). One issue with this possibility is that the kinesin stalk is on the order of tens of nm; therefore it cannot account for the several μm measured for <*d*>.

A second possible explanation is a change of <*d*> while gliding. The measured <*d*> is not influenced by the increases in the kinesin detachment rate, *k*_off_, under high ATP concentrations (*k*_off_ = 0.66 s^−1^ at 1 mM ATP to 0.0009 s^−1^ without ATP).^66^ However, the calculated <*d*> includes the increase of *k*_off_ by increased ATP concentration. Therefore, kinesin could easily detached from the MTs, resulting in larger apparent <*d*>.

A third explanation may be due to the deviation of the MT trajectories from the approximation model. The model is derived by assuming the infinitesimal deformation theory is applicable.^67^ When many kinesins are stretched, or detached from leading tips, they experience a large deformation and the model is no longer applicable at the microscopic level. To justify the approximation model, we estimated the maximum deflection at the free end, *y*_max_, by

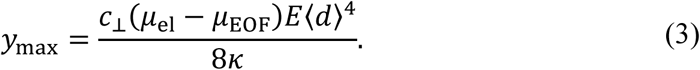

The ratio of *y*_max_ to <*d*> was smaller than 5%—2.8% for MT-2, 3.3% for MT-5, and 3.8% for MT-5’. The squared *y*_max_/<*d*> were up to 0.005, which satisfied the assumption of the infinitesimal deformation theory, i.e. squared *y*_max_/<*d*> should be much smaller than 1. Therefore, the approximation model was appropriate and we can reject the third reason.

By considering these explanations, we propose that the calculated <*d*> value should be regarded as an effective <*d*>, which differs from the actual distance between kinesins. Therefore, we proposed a strategy to directly measure *R*_MT_ in a flow cell to define the dimension of a microfluidic device rather than to calculate *R*_MT_ via *L*_p_ and <*d*> measured as the distance between kinesins. Once the effective <*d*> was calculated from *L*_p_ and the measured *R*_MT_, one can use the calculated <*d*> for any experimental setup to estimate *R*_MT_ for the design of a microfluidic device. As far as the cantilever model can be applied and equation (2) is effective, our design methodology requires an assay to measure *L*_p_ and RMT for deriving <*d*>, which can be applied to design the molecular sorting device.

In summary, integration of the bottom-up design of MT properties with the top-down design of a microfluidic device enabled us to create a highly efficient autonomous MT sorting system. For the bottom-up approach, the stiffness of MTs was modified by polymerizing conditions: the growth rate, nucleotides, and tau binding, and the relationship between their persistence length, *L*_p_, and radius of curvature, *R*_MT_, was determined. Stiff MTs showed larger *R*_MT_ than soft MTs as estimated by the cantilever model. Additional electrical modifications to soft MTs further decreased the *R*_MT_. For the top-down approach, the structure of the device and field intensity were optimized based on the measured *R*_MT_. Thus, the sorting efficiency reached ∼60% with *L*_p_-modified MTs and ∼80% with both surface charge density-and *L*_p_-modified MTs. The combined top-down and bottom-up design methodology is essential to attain autonomous directional control of molecular shuttles. In conjunction with other molecular shuttle-based functions such as transport, concentration, and detection, sorting functions in μTAS could be replaced by molecular shuttles.

## Methods

### Reagent preparation

Reagents were purchased from Sigma (St. Louis, MO) unless otherwise stated. Tubulin was purified from porcine brains by two cycles of polymerization and depolymerization followed by a phosphocellulose column as previously described.^68^ Recycled tubulin was purified from the tubulin by one more cycle of polymerization and depolymerization to remove any non-polymerizable tubulin. Succinimidyl ester-conjugated tetramethylrhodamine (C-1171; Invitrogen, Carlsbad, CA) or Alexa Fluor 488 (A-20000; Invitrogen) were labelled by adding 50-molar or 15- molar excess dye to tubulin, respectively.^69^ Human kinesin (amino acid residues 1–573) with an N-terminal histidine tag was purified as previously reported.^70^ Purified proteins were stored in liquid nitrogen. DNA molecules were purchased from JBioS (Saitama, Japan). These were hybridized by incubating 5’-biotinylated single strand DNA (ssDNA) with 5’-Alexa Fluor 488-labelled complementary ssDNA at 1:1 molar ratio at 37°C for >20 min. The sequence was 5’-GAG GTC TTA ACG GTG GAG GAT GGG GGT TAG TCC GGG GCG CAG ATT CGA AT-3’. We used BRB80 buffer (80 mM PIPES, 1 mM MgCl_2_, and 1 mM EGTA, pH 6.8 with KOH) for the dilution and suspension of proteins unless otherwise stated. Tau proteins (2N4R) were purchased from rPeptide (Athens, GA).

### Measurement of MT growth rate

The growth rate was measured for the elongated segment from a seed MT. Seed MTs were polymerized by incubating recycled tubulin and Alexa Fluor 488-labelled tubulin at 2:1 in the presence of 1 mM DTT (048-29224; Wako, Osaka, Japan) and 1 mM GMPCPP (NU-405S; Jena Bioscience, Jena, Germany) at 37°C for 30 min. The seeds were stabilized in 20 μM paclitaxel after polymerization and elongated from both ends by incubating with tubulin at a total concentration of 10 or 30 μM, which consisted of non-labelled tubulin, recycled tubulin, and tetramethylrhodamine-labelled tubulin at a molar ratio of 1.3:0.4:1, in the presence of 1 mM MgSO_4_ (131-00405; Wako) and 1 mM GTP at 37°C for 60 min. The final paclitaxel concentration during elongation was 2 μM. MTs were sampled at 0, 0.5, 1, 1.5, 2, 2.5, 3, 4, 5, 7.5, 10, 15, 20, 30, and 60 min after starting the incubation to measure the growth rate. Sampled MTs were diluted to 0.1 μM (below the critical concentration) in the presence of 20 μM paclitaxel, and centrifuged (163,000 × g and 27°C for 20 min) to remove the non-polymerized tubulin. The precipitated MTs were resuspended and introduced into a flow chamber constructed by two coverslips (C218181 and C024361; Matsunami Glass, Osaka, Japan) and 50μm thickness double-sided tape (400P50; Kyodo Giken Chemical, Saitama, Japan). After a 5 min incubation for non-specific binding of MTs to the glass substrate, the chamber was washed out and sealed with clear nail polish. The elongated length of the tetramethylrhodamine-labelled MT segment was measured with FIESTA by fitting the MT shape with the sub-pixel resolution via Gaussian fitting. Extremely short MTs (<500 nm), i.e., just after the start of elongation, could not be detected by FIESTA. When the number of undetectable MTs exceeded that of the detectable MTs, we did not use the image to measure MT length. The MT growth rate was defined by fitting initial time data to a linear equation where the slope reports the rate of growth (*R*^2^ > 0.90). We used only length distributions that were normally distributed.

### Experimental procedures for *L*_p_ measurement

MT-1 to MT-5 were extended from Alexa Fluor 488-labelled biotinylated MT seeds. Following the preparation of MT seeds as stated in the section above, they were biotinylated by incubating with a 20-fold molar excess of biotin-XX succinimidyl ester (B1606; Invitrogen) to tubulin at 37°C for 30 min. After quenching the unreacted biotin with a 200-fold molar excess of K-glutamate to tubulin at 37°C for 10 min, they were shortened by shearing through a 30-G syringe needle (90030; Osaka Chemical, Osaka, Japan).^5^ Seeds were stabilized with 20 μM paclitaxel after centrifugation, and elongated from both ends at 37°C for >20 min under five different conditions: GMPCPP and 10 μM tubulin with tau (MT-1) and without tau (MT-2), GTP and 10 μM tubulin with tau (MT-3) and without tau (MT-4), and GTP and 30 μM tubulin without tau (MT-5). The tau concentration was 1 μM, 10-fold lower than tubulin. All MTs were stabilized by 20 μM paclitaxel after polymerization.

Glass coverslips were cleaned in acetone, isopropanol, and HNO_3_ in order, and rinsed in deionized water (DIW). They were dried with nitrogen gas, and exposed to air plasma (Covance MP; Femto Science, Gyeonggi-Do, Korea). Then, they were immersed in a mixture of 1 mg ml^−1^ biotin-PEG-silane (MW3400, Biotin-PEG-SIL-3400-1g; Laysan Bio, Arab, AL), 30 mM HCl, and 97% ethanol in a nitrogen chamber overnight. They were rinsed in ethanol and DIW, and dried for storage at 4°C until their use.^32^

Flow chambers were constructed by bonding the biotin-coated coverslip to a non-coated coverslip with double-sided tape (10μm thickness, 7070W; Teraoka Seisakusho, Tokyo, Japan). We introduced 2 mg ml^−1^ streptavidin (192-11644; Wako) and incubated for 3 min. After washing out the chambers with BRB80, the partially biotinylated MTs (MT-1 to MT-5) were immobilized via biotin-streptavidin bindings by a 5 min incubation. Non-immobilized MTs were removed by flowing through BRB80, which included an O2 scavenger system (BRB8O-O2; 8.0 μg ml^−1^ catalase, 25 mM D-glucose, 20 μg ml^−1^ glucose oxidase, 1% β-ME, 20 mM DTT, 3.0 mM 1,1’-ferrocenedimethanol (322-49071; Wako), and 20 μM paclitaxel in BRB80). Finally, the chamber was sealed with clear nail polish to prevent flow.

*L*_p_ was measured at room temperature (~27°C). MT shapes were analysed with a custom-written MATLAB algorithm (MathWorks, Natick, MA) as reported previously.^60^ Briefly, we expressed the MT shape as a superposition of Fourier modes and measured MT bending energy from their deformation on the assumption that the elongated segment behaved as a cantilever clamped at an immobilized biotinylated segment. Since MT fluctuations originated only from thermal energy in the sealed chamber, the MT bending energy was equated with thermal energy, which enabled us to derive *L*_p_. The analysis procedure has been published in previous reports (see Supporting Information and Supplementary Fig. 2 for details).^27,34,60^

### Fabrication of the PDMS device

A bare silicon wafer was dehydrated at 200°C for 5 min and spincoated with hexamethylsilazane at 500 rpm for 5 s and 3,000 rpm for 40 s to increase the adhesiveness of an SU-8 resist (1H-D7; Mikasa, Tokyo, Japan). After baking the hexamethylsilazane at 120°C for 5 min, an SU-8 3010 photoresist (MicroChem, Westborough, MA) was spincoated at 500 rpm for 10 s (ramp: 100 rpm s^−1^) and 3,000 rpm for 30 s (ramp: 300 rpm s^−1^), followed by baking at 65°C for 2 min, 95°C for 3 min, and 65°C for 1 min in that order. The channel pattern drawn on a Cr mask (HS Hardmask Blanks; Clean Surface Technology Co., Samukawa, Japan) was transferred by a mask aligner (PEM800; Union, Tokyo, Japan) with an exposure energy of 200 mJ cm^−2^ and baked at 65°C for 1 min, 95°C for 2 min, and 65°C for 3 min. The wafer was cooled at room temperature for 5-10 min and SU-8 was developed by immersion in a developer (Microchem) at 40°C for 3 min. After rinsing with isopropyl alcohol at 40°C for 10 s and drying, the SU-8 mold was silanized overnight in a vacuum chamber filled with gaseous trichloro(1H, 1H,2H,2H-perfluorooctyl) silane to enable easy peeling of the cured PDMS.

A PDMS prepolymer was mixed with a curing agent (SILPOT 184 W/C; Dow Corning Toray, Tokyo, Japan) at a ratio of 10:1 (w/w) and cast onto the SU-8 mold at a thickness of ~5 mm. It was then degassed in a vacuum chamber for 30 min and cured at 80°C for 2 h. The cured PDMS was peeled from the mold and punched with a biopsy punch (Sterile Dermal Biopsy Punch, 3 mm; Kai Industries, Tokyo, Japan) to make reservoirs. Coverslips were cleaned in 10 N KOH solution at room temperature for 24 h and rinsed twice by ultrasonication in DIW at room temperature for 20 min. They were then immersed in 20% ethanol solution at room temperature for 10 min and rinsed in DIW, followed by drying with nitrogen gas. The PDMS and the cleaned coverslips were exposed to air plasma and bonded permanently.

### Experimental procedures for *R*_MT_ measurement

MT-1, MT-2, and MT-5 without GMPCPP seeds were polymerized for *R*_MT_ measurement. MT-1 was polymerized under 1 mM GMPCPP and 1 mM tau using 10 μM tubulin. MT-2 was polymerized under 1 mM GMPCPP and 10 μM tubulin without tau. MT-5 was polymerized under 1 mM GTP and 30 μμ tubulin without tau. For the preparation of MT-5’, the shortened MT-5 was biotinylated and elongated only from the plus end by incubating it with a mixture of non-fluorophore-labelled, N-ethylmaleimide (NEM)-treated, and fluorophore-labelled tubulin in the presence of 1 mM MgSO_4_ and 1 mM GTP at 37°C for 30 min. NEM-treated tubulin was prepared by incubating recycled tubulin with 1 mM GTP and 300-500 molar excess of NEM (10 mM) at 4°C for 10 min, followed by incubation with 0.56% β-ME at 4°C for 10 min to inactivate the excess NEM.^69^ MT-5 biotinylated at the minus-end was centrifuged, resuspended with 20 μM paclitaxel, and labelled with the hybridized DNA at 37°C.

A PDMS channel was coated with 2 mg ml^−1^ Pluronic F108 (BASF, Ludwigshafen, Germany) to prevent kinesin binding onto the PDMS surfaces, and then 0.08 mg ml^−1^ kinesin with 1.9 mg ml^−1^ casein was immobilized. These solutions introduced from reservoir A were incubated for 5 min and washed out with BRB80. Four types of MTs were introduced and all reservoirs were filled with 0.5 mM ATP diluted with BRB80-O2 and platinum electrodes were inserted into reservoirs A and B. An electric field from reservoir B to A was applied with an average intensity of 5.3 kV m^−1^ (E3612; Agilent Technologies, Tokyo, Japan).

MTs were tracked at the leading tips with Mark2 software (provided by Dr. Kenya Furuta, National Institute of Information and Communications Technology, Kobe, Japan), and their trajectories were fitted by equation (1) to obtain RMT. MTs gliding discontinuously or turning on a pivot were removed from the analysis.

### MT sorting

The PDMS channel was coated with Pluronic F108, kinesin, and casein sequentially. To immobilize MTs on the landing area surface only, 5 μL of BRB80-O2 was introduced to reservoirs A and B, and then 5 μL of an MT mixture (MT-2 and MT-5, or MT-2 and MT-5’) was loaded into reservoir C, resulting in a flow toward empty reservoir D from the other reservoirs. The MT solution in reservoir C was removed and the landing area was washed out by 5μL BRB80-O_2_. An electric field was applied from reservoir B to A via platinum electrodes inserted into ATP-filled reservoirs. The average field intensity was set to 3.0 kV m^-1^. MT sorting was visualized with ImageJ software (National Institutes of Health, Bethesda, MD) with the MTrackJ plug-in.^71^ MTs were tracked at the leading tips at 15-s intervals once they entered from MT alignment to the sorting area along the *y*-axis.

### Optical imaging and analysis

MT and DNA molecules were observed under an IX73 inverted epifluorescence microscope (Olympus, Tokyo, Japan) with an excitation filter (GFP/DsRed-A-OMF; Opto-Line International, Inc., Wilmington, MA), a complementary metal-oxide-semiconductor camera (ORCA-Flash 4.0 V2; Hamamatsu Photonics, Hamamatsu, Japan), image splitting optics (W-VIEW GEMINI; Hamamatsu Photonics) with a bandpass emitter (FF01-512/25-25 and FF01-630/92-25; Semrock, Rochester, NY) and dichroic mirror (FF560-FDi01-25x36; Semrock), and oil-immersion objectives. The magnification, exposure time, frame rate, and recording period were 100 × (NA 1.4), 100 ms, 2.5 frame s^−1^, and 200 s for observing MT thermal fluctuation, and 60 × (NA 1.35), 200 ms, 0.33-0.5 frame s^−1^, and >1 h for observing MT gliding, respectively. An ND6 filter was used with a shutter (VMM-D3; Uniblitz, Rochester, NY). Optical images were stored as sequential image files in TIFF format using HCImage software (Hamamatsu Photonics).

Multiple significance tests were performed among all data by Steel-Dwass tests at a critical value of *p* < 0.01, and normality or log-normality was tested among outlier-removed data by Lilliefors tests at a critical value of *p* > 0.05. All curve fittings were performed by the least squares methods of MATLAB.

## Acknowledgements

This study was partially supported by Precursory Research for Embryonic Science and Technology (PRESTO) from the Japan Science and Technology Agency (JST); Japan Society for the Promotion of Science (JSPS) KAKENHI Grant Number 25709018; Kyoto University Supporting Program for Interaction-Based Initiative Team Studies (SPIRITS) as part of the Program for Promoting the Enhancement of Research Universities, the Ministry of Education, Culture, Sports, Science and Technology (MEXT), Japan; Kyoto University Nano Technology Hub in “Nanotechnology Platform Project” sponsored by MEXT, Japan. N.I. was supported by Grant-in-Aid for JSPS Research Fellow Grant Number 262439. J.L.R. is supported by a grant from the Mathers Foundation, Moore Foundation grant 4308.1, National Science Foundation INSPIRE award MCB-1344203, and the DoD ARO MURI 67455-CH-MUR MURI award. T.L.H. was supported by Wisconsin Space Grant and UWL Faculty Research Grant. We also thank Dr. K. Furuta for the software, Mark2, for tracking the MT trajectories.

## Author Contributions

N.I. and R.Y. designed the experiments. N.I. performed the experiments and analysed the data. N.I., H.S., H.K., T.L.H., J.L.R., and R.Y. discussed and interpreted the results. N.I., T.L.H., J.L.R., and R.Y. wrote the paper. All authors have approved the final version of the manuscript.

## Additional information

**Competing financial interests:** The authors declare no competing financial interests.

## References

1. Verhey, K. J. & Hammond, J. W. Traffic control: regulation of kinesin motors. Nat. Rev. Mol. Cell Biol. 10, 765–777 (2009).

2. Schnitzer, M. J. & Block, S. M. Kinesin hydrolyses one ATP per 8-nm step. Nature 388, 386–390 (1997).

3. Vale, R. D., Schnapp, B. J., Reese, T. S. & Sheetz, M. P. Organelle, bead, and microtubule translocations promoted by soluble factors from the squid giant axon. Cell 40, 559–569 (1985).

4. Agarwal, A. & Hess, H. Biomolecular motors at the intersection of nanotechnology and polymer science. Prog. Polym. Sci. 35, 252–277 (2010).

5. Lin, C. T., Kao, M. T., Kurabayashi, K. & Meyhofer, E. Self-contained, biomolecular motor-driven protein sorting and concentrating in an ultrasensitive microfluidic chip. Nano Lett. 8, 1041–1046 (2008).

6. Steuerwald, D., Fruh, S. M., Griss, R., Lovchik, R. D. & Vogel, V. Nanoshuttles propelled by motor proteins sequentially assemble molecular cargo in a microfluidic device. Lab Chip 14 (19) 3729–3738 (2014).

7. Ramachandran, S., Ernst, K. H., Bachand, G. D., Vogel, V. & Hess, H. Selective loading of kinesin-powered molecular shuttles with protein cargo and its application to biosensing. Small 2, 330–334 (2006).

8. Fischer, T., Agarwal, A. & Hess, H. A smart dust biosensor powered by kinesin motors. Nat. Nanotechnol. 4, 162–166 (2009).

9. Taira, S. et al. Selective detection and transport of fully matched DNA by DNA-loaded microtubule and kinesin motor protein. Biotechnol. Bioeng. 95, 533–538 (2006).

10. Hiyama, S., Moritani, Y., Gojo, R., Takeuchi, S. & Sutoh, K. Biomolecular-motor-based autonomous delivery of lipid vesicles as nano-or microscale reactors on a chip. Lab Chip 10, 2741–2748 (2010).

11. Hiratsuka, Y., Tada, T., Oiwa, K., Kanayama, T. & Uyeda, T. Q. P. Controlling the direction of kinesin-driven microtubule movements along microlithographic tracks. Biophys. J. 81, 1555–1561 (2001).

12. Clemmens, J., Hess, H., Howard, J. & Vogel, V. Analysis of microtubule guidance in open microfabricated channels coated with the motor protein kinesin. Langmuir 19, 1738–1744 (2003).

13. Kim, T., Meyhofer, E. & Hasselbrink, E. F. Biomolecular motor-driven microtubule translocation in the presence of shear flow: modeling microtubule deflection due to shear. Biomed. Microdevices 9, 501–511 (2007).

14. Agayan, R. R. et al. Optimization of isopolar microtubule arrays. Langmuir 29, 2265–2272 (2013).

15. Hutchins, B. M., Platt, M., Hancock, W. O. & Williams, M. E. Directing transport of CoFe2O4-functionalized microtubules with magnetic fields. Small 3, 126–131 (2007).

16. van den Heuvel, M. G., de Graaff, M. P. & Dekker, C. Molecular sorting by electrical steering of microtubules in kinesin-coated channels. Science 312, 910-914 (2006).

17. van den Heuvel, M. G., de Graaff, M. P. & Dekker, C. Microtubule curvatures under perpendicular electric forces reveal a low persistence length. Proc. Natl. Acad. Sci. U. S. A. 105, 7941–7946 (2008).

18. Kim, T., Kao, M. T., Hasselbrink, E. F. & Meyhofer, E. Active alignment of microtubules with electric fields. Nano Lett. 7, 211–217 (2007).

19. Isozaki, N. et al. Control of microtubule trajectory within an electric field by altering surface charge density. Sci. Rep. 5, 7669 (2015).

20. Cassimeris, L., Gard, D., Tran, P. T. & Erickson, H. P. XMAP215 is a long thin molecule that does not increase microtubule stiffness. J. Cell Sci. 114, 3025–3033 (2001).

21. Hawkins, T., Mirigian, M., Selcuk Yasar, M. & Ross, J. L. Mechanics of microtubules. J. Biomech. 43, 23–30 (2010).

22. Mizushima-Sugano, J., Maeda, T. & Miki-Noumura, T. Flexural rigidity of singlet microtubules estimated from statistical analysis of their contour lengths and end-to-end distances. BBA - Gen. Subjects 755, 257–262 (1983).

23. Dye, R. B., Fink, S. P. & Williams, R. C. Taxol-induced flexibility of microtubules and its reversal by MAP-2 and Tau. J. Biol. Chem. 268, 6847–6850 (1993).

24. Mickey, B. & Howard, J. Rigidity of Microtubules Is Increased by Stabilizing Agents. J. Cell Biol. 130, 909–917 (1995).

25. Felgner, H. et al. Domains of neuronal microtubule-associated proteins and flexural rigidity of microtubules. J. Cell Biol. 138, 1067–1075 (1997).

26. Kis, A. et al. Nanomechanics of Microtubules. Phys. Rev. Lett. 89, 248101 (2002).

27. Janson, M. E. & Dogterom, M. A bending mode analysis for growing microtubules: evidence for a velocity-dependent rigidity. Biophys. J. 87, 2723-2736 (2004).

28. Brangwynne, C. P. et al. Bending Dynamics of Fluctuating Biopolymers Probed by Automated High-Resolution Tracking. Biophys. J. 93, 346–359 (2007).

29. van Mameren, J., Vermeulen, K. C., Gittes, F. & Schmidt, C. F. Leveraging Single Protein Polymers To Measure Flexural Rigidity. J. Phys. Chem. B 113, 3837–3844 (2009).

30. Kawaguchi, K. & Yamaguchi, A. Temperature dependence rigidity of non-taxol stabilized single microtubules. Biochem. Biophys. Res. Commun. 402, 66–69 (2010).

31. Hawkins, T. L., Sept, D., Mogessie, B., Straube, A. & Ross, J. L. Mechanical properties of doubly stabilized microtubule filaments. Biophys. J. 104, 1517–1528 (2013).

32. Portran, D. et al. MAP65/Ase1 promote microtubule flexibility. Mol. Biol. Cell 24, 1964–1973 (2013).

33. Bouxsein, N. F. & Bachand, G. D. Single Filament Behavior of Microtubules in the Presence of Added Divalent Counterions. Biomacromolecules 15, 3696–3705 (2014).

34. Gittes, F., Mickey, B., Nettleton, J. & Howard, J. Flexural rigidity of microtubules and actin filaments measured from thermal fluctuations in shape. J. Cell Biol. 120, 923–934 (1993).

35. Kikumoto, M., Kurachi, M., Tosa, V. & Tashiro, H. Flexural rigidity of individual microtubules measured by a buckling force with optical traps. Biophys. J. 90, 1687–1696 (2006).

36. Pampaloni, F. et al. Thermal fluctuations of grafted microtubules provide evidence of a length-dependent persistence length. Proc. Natl. Acad. Sci. U.S.A 103, 10248–10253 (2006).

37. Kurachi, M., Hoshi, M. & Tashiro, H. Buckling of a single microtubule by optical trapping forces: Direct measurement of microtubule rigidity. Cell Motil. Cytoskeleton 30, 221–228 (1995).

38. Taute, K. M., Pampaloni, F., Frey, E. & Florin, E.-L. Microtubule Dynamics Depart from the Wormlike Chain Model. Phys. Rev. Lett. 100, 028102 (2008).

39. Mitchison, T. & Kirschner, M. Dynamic instability of microtubule growth. Nature 312, 237–242 (1984).

40. Zanic, M., Widlund, P. O., Hyman, A. A. & Howard, J. Synergy between XMAP215 and EB1 increases microtubule growth rates to physiological levels. Nat. Cell Biol. 15, 688–693 (2013).

41. Kerssemakers, J. W. J. et al. Assembly dynamics of microtubules at molecular resolution. Nature 442, 709–712 (2006).

42. Dogterom, M. & Yurke, B. Measurement of the Force-Velocity Relation for Growing Microtubules. Science 278, 856–860 (1997).

43. Kinoshita, K., Arnal, I., Desai, A., Drechsel, D. N. & Hyman, A. A. Reconstitution of Physiological Microtubule Dynamics Using Purified Components. Science 294, 1340–1343 (2001).

44. Vasquez, R. J., Howell, B., Yvon, A. M., Wadsworth, P. & Cassimeris, L. Nanomolar concentrations of nocodazole alter microtubule dynamic instability in vivo and in vitro. Mol. Biol. Cell 8, 973–985 (1997).

45. Komarova, Y. et al. Mammalian end binding proteins control persistent microtubule growth. J. Cell Biol. 184, 691–706 (2009).

46. Ruhnow, F., Zwicker, D. & Diez, S. Tracking Single Particles and Elongated Filaments with Nanometer Precision. Biophys. J. 100, 2820–2828 (2011).

47. Korten, T. & Diez, S. Setting up roadblocks for kinesin-1: mechanism for the selective speed control of cargo carrying microtubules. Lab Chip 8, 1441–1447 (2006).

48. Morshed, B., Shams, M. & Mussivand, T. Analysis of Electric Fields inside Microchannels and Single Cell Electrical Lysis with a Microfluidic Device. Micromachines 4, 243 (2013).

49. Janson, M. E. & Dogterom, M. Scaling of Microtubule Force-Velocity Curves Obtained at Different Tubulin Concentrations. Phys. Rev. Lett. 92, 248101 (2004).

50. Schaedel, L. et al. Microtubules self-repair in response to mechanical stress. Nat. Mater. 14, 1156–1163 (2015).

51. Wada, S. et al. Effect of length and rigidity of microtubules on the size of ring-shaped assemblies obtained through active self-organization. Soft Matter 11, 1151–1157 (2015).

52. Vale, R. D., Coppin, C. M., Malik, F., Kull, F. J. & Milligan, R. A. Tubulin GTP hydrolysis influences the structure, mechanical properties, and kinesin-driven transport of microtubules. J. Biol. Chem. 269, 23769–23775 (1994).

53. Lopez, B. J. & Valentine, M. T. Mechanical effects of EB1 on microtubules depend on GTP hydrolysis state and presence of paclitaxel. Cytoskeleton 71, 530-541 (2014).

54. Yajima, H. et al. Conformational changes in tubulin in GMPCPP and GDP-taxol microtubules observed by cryoelectron microscopy. J. Cell Biol. 198, 315–322 (2012).

55. Müller-Reichert, T., Chrétien, D., Severin, F. & Hyman, A. A. Structural changes at microtubule ends accompanying GTP hydrolysis: Information from a slowly hydrolyzable analogue of GTP, guanylyl (α, β)methylenediphosphonate. Proc. Natl. Acad. Sci. U.S.A. 95, 3661–3666 (1998).

56. Elie-Caille, C. et al. Straight GDP-Tubulin Protofilaments Form in the Presence of Taxol. Curr. Biol. 17, 1765–1770 (2007).

57. Kadavath, H. et al. Tau stabilizes microtubules by binding at the interface between tubulin heterodimers. Proc. Natl. Acad. Sci. U.S.A 112, 7501–7506 (2015).

58. Kasas, S. et al. Mechanical Properties of Microtubules Explored Using the Finite Elements Method. Chemphyschem 5, 252–257 (2004).

59. Felgner, H., Frank, R. & Schliwa, M. Flexural rigidity of microtubules measured with the use of optical tweezers. J. Cell Sci. 109, 509–516 (1996).

60. Hawkins, T. et al. Perturbations in Microtubule Mechanics from Tubulin Preparation. Cel. Mol. Bioeng. 5, 227–238 (2012).

61. Venier, P., Maggs, A. C., Carlier, M. F. & Pantaloni, D. Analysis of Microtubule Rigidity Using Hydrodynamic Flow and Thermal Fluctuations. J. Biol. Chem. 269, 13353–13360 (1994).

62. Zhang, J. & Wang, C. Molecular structural mechanics model for the mechanical properties of microtubules. Biomech. Model. Mechanobiol. 13, 1175–1184 (2014).

63. Ikuta, J. et al. Tug-of-war of microtubule filaments at the boundary of a kinesin-and dynein-patterned surface. Sci. Rep. 4, 5281 (2014).

64. Fallesen, T. L., Macosko, J. C. & Holzwarth, G. Force-velocity relationship for multiple kinesin motors pulling a magnetic bead. Eur. Biophys. J. 40, 1071–1079 (2011).

65. Fallesen, T. L., Macosko, J. C. & Holzwarth, G. Measuring the number and spacing of molecular motors propelling a gliding microtubule. Phys. Rev. E Stat. Nonlin. Soft Matter Phys. 83, 011918 (2011).

66. Hancock, W. O. & Howard, J. Kinesin’s processivity results from mechanical and chemical coordination between the ATP hydrolysis cycles of the two motor domains. Proc. Natl. Acad. Sci. U.S.A. 96, 13147–13152 (1999).

67. Timoshenko, S. P. Strength of Materials. 3rd edn, Vol. pt. 1: Elementary theory and problems (Van Nostrand Reinhold, New York, NY, 1958).

68. Williams, R. C., Jr. & Lee, J. C. Preparation of tubulin from brain. Methods Enzymol. **85 Pt B,** 376–385 (1982).

69. Hyman, A. et al. Preparation of modified tubulins. Methods Enzymol. 196, 478-485 (1991).

70. Yokokawa, R., Tarhan, M. C., Kon, T. & Fujita, H. Simultaneous and bidirectional transport of kinesin-coated microspheres and dynein-coated microspheres on polarity-oriented microtubules. Biotechnol. Bioeng. 101, 1–8 (2008).

71. Meijering, E., Dzyubachyk, O. & Smal, I. in Methods Enzymol. Vol. 504 (ed P. M. Conn) Ch. 9, 183–200 (Academic Press, 2012).

